## Supplementary Material for "Chromosome organization shapes replisome dynamics in *Caulobacter crescentus*"

### Affiliations:

### Content:

**Supplementary methods**

- Plasmid and strain construction
- Microplate reader experiments

**Supplementary table and figures**

**Supplementary references**

### Supplementary methods

#### Plasmid and strain construction

The integrative plasmid *pDnaN-sfGFP* was first constructed by replacing the *mCherry* into *sfGFP* in the plasmid of pDnaN-RFP, and was introduced into *C. crescentus* cells by conjugation with the *E. coli* strain S17-1[1]. Notably, we tried the electroporation protocol but failed.

The integrative plasmid $P_{xyl}$*-zapT-mScarlet-I* was constructed based on the previously published $P_{xyl}$*-ftsZ-sfGFP* plasmid[2], where *mScarlet-I* was tagged to the C-terminal of full-length *zapT* gene and then replaced the *ftsZ-sfGFP*. The $P_{xyl}$*-zapT-mScarlet-I* was then introduced into *C. crescentus* cells by electroporation. The integrative plasmid $P_{xyl}$*-rsaA* was constructed by amplifying *rsaA* gene (from promoter to coding sequence) using the same primers as reported previously[3], and then replacing *ftsZ-sfGFP* gene in the $P_{xyl}$*-ftsZ-sfGFP* plasmid. The $P_{xyl}$*-rsaA* was introduced into *C. crescentus* cells by electroporation.

The integrative plasmid harboring orthogonal ParB/*parS* system was constructed by tagging the cassette gene of "mCherry-ParB-*parS* (P1)" or "GFP-ParB-*parS* (pMT1)" to ~1000 bp of chromosome sequence from the left or right arm respectively, followed by the insertion into pMCS-2 or pMCS-1[4]. Cassette genes were from pWX995 and pWX967. Notably, the site-specific ~1000 bp of chromosome sequence was carefully chosen to make sure that the integrated ParB/*parS* constructs did not affecting the expression of essential genes in *C. crescentus*[5]. The ultimate plasmids for genomic integration on left and right arms were sequentially transformed into *C. crescentus* by electroporation or transferred from one strain to another by phage Cr30- mediated transduction.

To construct the *parAK20R* mutant, the cassette encoding such ParA point-mutant was transduced into wide-type *C. crescentus* at the $P_{xyl}$ region by phage transduction. To induce the ParA mutant, 0.2% wt/vol xylose was added in cell cultures 2 h before imaging. Strains including Δ*smc*, *flip1-5*, Δ*rsaA* were from previous studies[3,6]. All details of plasmids and strains used are listed in **Table S1**.

#### Microplate reader experiments

Overnight cell culture of *C. crescentus* cells ($OD_{660}$ ~1) were dilute 1000 times into the fresh medium. A 200 μL of diluted culture was loaded into 96-well plate for measuring the growth curve. The microplate reader device (Thermo Scientific, Multiskan FC) was set on 28°C or 32°C under shaking conditions (200 rpm), and recorded the absorbance ($A_{660}$) every 20 min over 40 h. Five biological repeats were measured for each strain. To calculate the doubling time ($T_{db}$), the time (ΔT) for cells growing from early-log phase (OD1, $OD_{660}$ ~0.1) to mid-log phase (OD2, $OD_{660}$ ~0.3) was count, and use the formula: $T_{db} = \ln 2 / (\ln(OD2/OD1)) / \Delta T)$.

### Supplementary table and figures

**Table 1:** details of plasmids and strains used in this study

| Plasmids (antibiotic marker) | Descriptions | Sources (# lab stock) |
|---|---|---|
| *pDnaN-mCherry* (spc$^{R}$) | Integrative plasmid with fusion of sfGFP to the C-terminal of DnaN | Justine Collier lab[7] |
| *pDnaN-sfGFP* (spc$^{R}$) | Integrative plasmid with fusion of sfGFP to the C-terminal of DnaN | This study (#512) |
| *pWX967* (kan$^{R}$) | Background strain containing GFP-ParB$^{MT1}$-*parS*$^{MT1}$ cassette | Xindan Wang lab[8,9] |
| *pWX995* (spc$^{R}$) | Background strain containing mCherry-ParB$^{P1}$-*parS*$^{P1}$ cassette | |
| *pMCS-1* (spc$^{R}$) | Empty integrative plasmids | Martin Thanbichler lab[4] |
| *pMCS-2* (kan$^{R}$) | | |
| *pL1* (kan$^{R}$) | Integrative pMCS-2 plasmid containing mCherry-ParB$^{P1}$-*parS*$^{P1}$ ahead of the sequence (-3705700 bp to -3704514 bp) | This study (#559) |
| *pR1* (spc$^{R}$) | Integrative pMCS-1 plasmid containing GFP-ParB$^{MT1}$-*parS*$^{MT1}$ after the sequence (+336613 bp to +337800 bp) | This study (#564) |
| *pL5'* (kan$^{R}$) | Integrative pMCS-2 plasmid containing mCherry-ParB$^{P1}$-*parS*$^{P1}$ ahead of the sequence (-2357000 bp to -2358500 bp) | This study (#563) |
| *pR5'* (kan$^{R}$) | Integrative pMCS-1 plasmid containing GFP-ParB$^{MT1}$-*parS*$^{MT1}$ after the sequence (+1683240 bp to +1684700 bp) | This study (#568) |
| *Pxyl-ftsZ-sfGFP* | Integrative plasmid with fusion of sfGFP to the C-terminal of FtsZ under $P_{xyl}$ promoter | Lab stock (#500) |
| $P_{xyl}$-*zapT-mScarlet-I* (kan$^{R}$) | Integrative plasmid with fusion of mScarlet-I to the C-terminal of ZapT under $P_{xyl}$ promoter | This study (#522) |
| $P_{xyl}$-*rsaA* (kan$^{R}$) | Integrative plasmid with *rsaA* gene with its native promoter under *Pxyl* promoter | This study (#592) |
| **Strains (antibiotic marker)** | **Descriptions** | **Sources** |
| CB15N | Wild-type NA1000 *Caulobacter crescentus* (synchronizable) | Lab stock |
| CB15N *:: dnaN-mCherry* (spc$^{R}$) | *pDnaN-mCherry* integrated into CB15N at native *dnaN* site | Justine Collier lab[7] |
| CB15N *:: dnaN-sfGFP* (spc$^{R}$) | *pDnaN-sfGFP* integrated into CB15N at native *dnaN* site | This study |
| CB15N *:: dnaN-sfGFP* (spc$^{R}$) *::* $P_{xyl}$-*zapT-mScarlet-I* (kan$^{R}$) | Transformation of $P_{xyl}$-*zapT-mScarlet-I* into CB15N *:: dnaN-sfGFP* | This study |
| CB15N *:: parB-gfp* | CB15N containing *ParB-gfp* at its native site | Martin Thanbichler lab[10] |
| CB15N *:: parB-gfp :: dnaN-mCherry* (spc$^{R}$) | Transduction of *dnaN-mCherry* from CB15N *:: dnaN-mCherry* into CB15N *:: parB-gfp* | This study |
| CB15N :: $P_{lacI}$-*lacI (hfA locus) ::* $P_{lac}$-*dnaA (dnaA locus) :: ISce1@ori (tet*$^{R}$*) ::* $P_{xyl}$*_parAK20R (gent*$^{R}$*)* | The background strain containing $P_{xyl}$-*parAK20R* gene | Anjana Badrinarayanan lab[11] |

| | | |
|---|---|---|
| CB15N *:: parB-gfp :: dnaN-mCherry* (spc$^R$) *::$P_{xyl}$-parAK20R* (gen$^R$) | Transduction of *$P_{xyl}$-ParAK20R* from the background strain above into CB15N *:: parB-gfp :: dnaN-mCherry* | This study |
| CB15N *:: L1* (kan$^R$) | Integration of *pL1* into CB15N | This study |
| CB15N *:: R1* (spc$^R$) | Integration of *pR1* into CB15N | This study |
| CB15N *:: L1* (kan$^R$) *:: R1* (spc$^R$) | Transduction of *pL1* from CB15N *:: L1* into CB15N *:: R1* | This study |
| CB15N *:: L5'* (kan$^R$) | Integration of *pL5* into CB15N | This study |
| CB15N *:: R5'* (spc$^R$) | Integration of *pR5* into CB15N | This study |
| CB15N *:: L5'*(kan$^R$) *:: R5'* (spc$^R$) | Transduction of *pL5* from CB15N *:: L5* into CB15N *:: R5* | This study |
| CB15N Δ*smc* (kan$^R$) | *smc* deletion in CB15N | Tung Le lab[6] |
| CB15N *flip1-5* | CB15N with partial chromosome (+3611 to +4038 kb) inversed | |
| CB15N Δ*rsaA* | *rsaA* deletion in CB15N | |
| CB15N Δ*smc* (kan$^R$) *:: dnaN-sfGFP* (spc$^R$) | *pDnaN-sfGFP* integrated into CB15N Δ*smc* | This study |
| CB15N *flip1-5 :: dnaN-sfGFP* (spc$^R$) | *pDnaN-sfGFP* integrated into CB15N *flip1-5* | This study |
| CB15N Δ*rsaA :: dnaN-sfGFP* (spc$^R$) | *pDnaN-sfGFP* integrated into CB15N Δ*rsaA* | This study |
| CB15N Δ*rsaA :: dnaN-sfGFP* (spc$^R$) *:: $P_{xyl}$-rsaA* (kan$^R$) | Transformation of *$P_{xyl}$-rsaA* into CB15N Δ*rsaA :: dnaN-sfGFP* | This study |
| CB15N *:: dnaN-sfGFP* (spc$^R$) *:: $P_{xyl}$-rsaA* (kan$^R$) | Transformation of *$P_{xyl}$-rsaA* into CB15N *:: dnaN-sfGFP* | This study |
| CB15N *:: L2* | #102_tet_3645906 (CB15N targeted locus: -371033 bp) | Patrick Viollier lab[12] |
| CB15N *:: R2* | #85_tet_433392 (CB15N targeted locus: +433392 bp) | |
| CB15N *:: L3* | #11_tet_3029646 (CB15N targeted locus: -987304 bp) | |
| CB15N *:: R3* | #3_tet_957206 (CB15N targeted locus: +983237 bp) | |
| CB15N *:: L4* | #77_tet_2635677 (CB15N targeted locus: -1381361 bp) | |
| CB15N *:: R4* | #89_tet_1349907 (CB15N targeted locus: +1375883 bp) | |
| CB15N *:: L5* | #97_tet_2495294 (CB15N targeted locus: -1521654 bp) | |
| CB15N *:: R5* | #8_tet_1498826 (CB15N targeted locus: +1524802 bp) | |
| CB15N *:: L5* | #2_tet_2351851 (CB15N targeted locus: -1665102 bp) | |
| CB15N *:: R5* | #129_tet_1661370 (CB15N targeted locus: +1687346 bp) | |

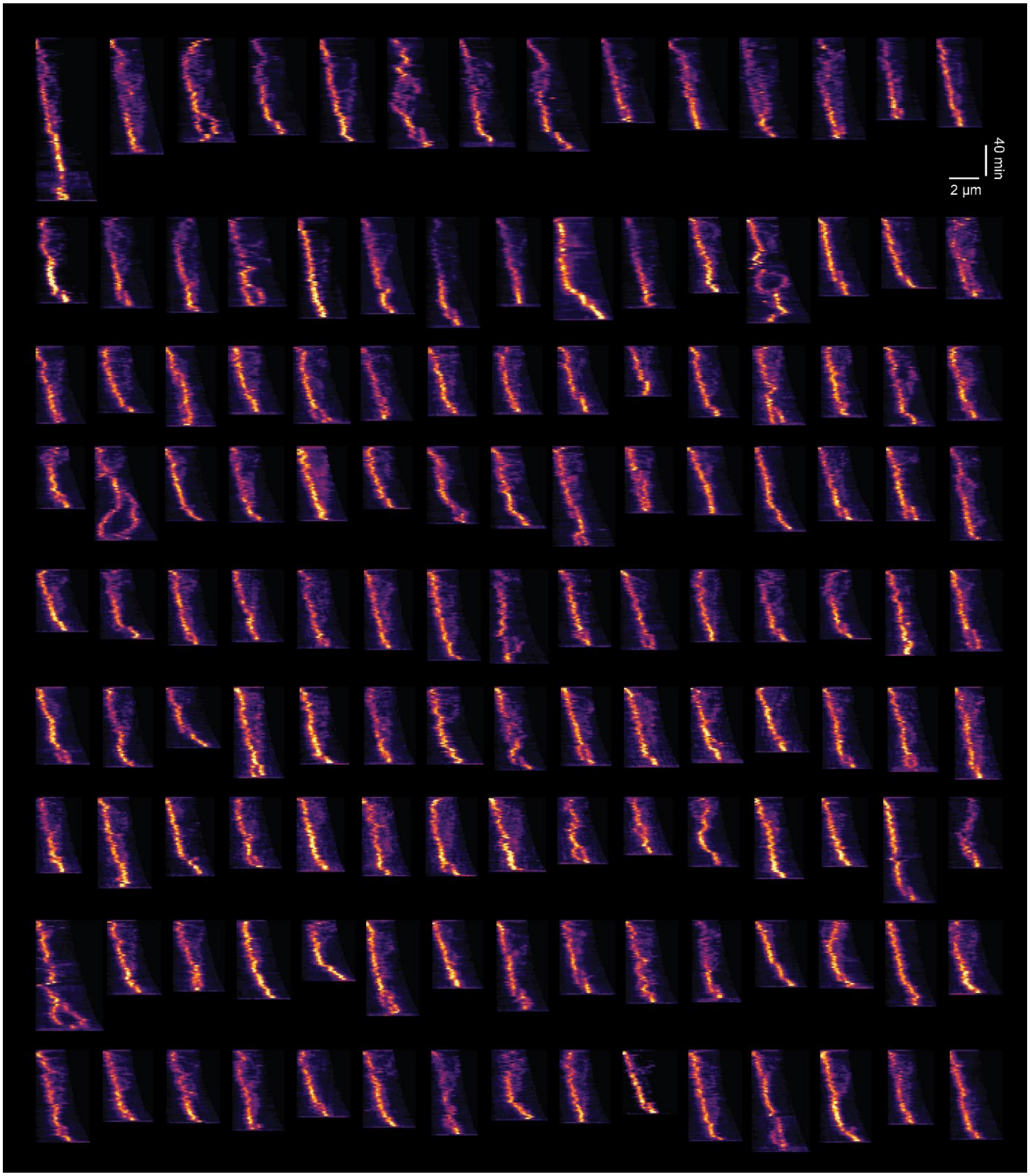


**Fig. S1**: Kymographs of time-lapse imaging of the CB15N*::dnaN-sfGFP* cells.

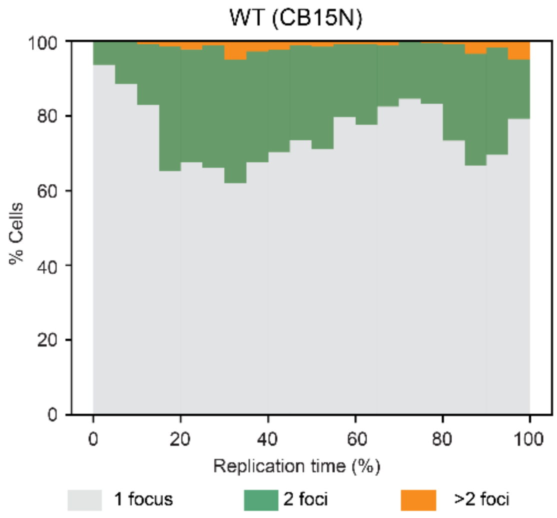


**Fig. S2:** Distribution of CB15N::*dnaN-sfGFP* cells that contain 1, 2, and >2 detected DnaN foci.

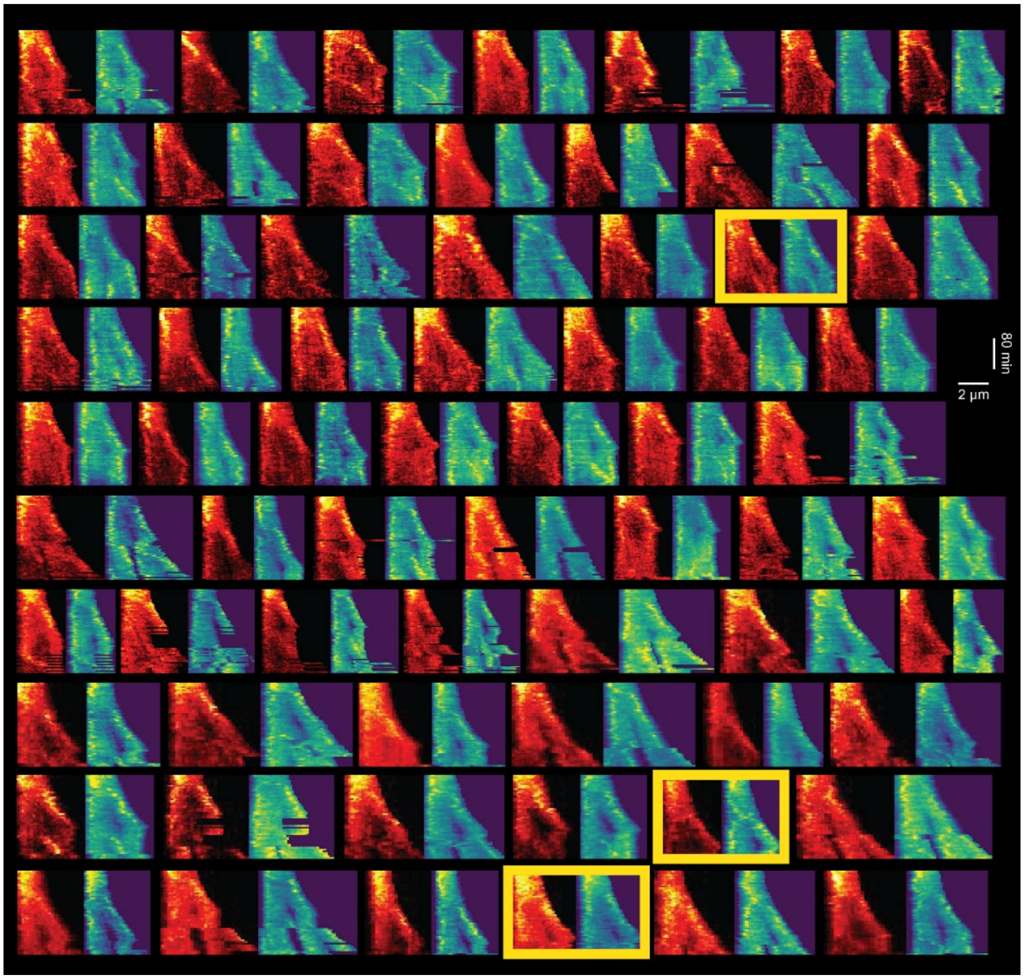


**Fig. S3:** Kymographs of L1 (left, pseudo-colored in Red-Hot) and R1 (right, pseudo-colored in MPI-viridis) fluorescence. Three cells highlighted in yellow boxes are considered to have different loci splitting sites.

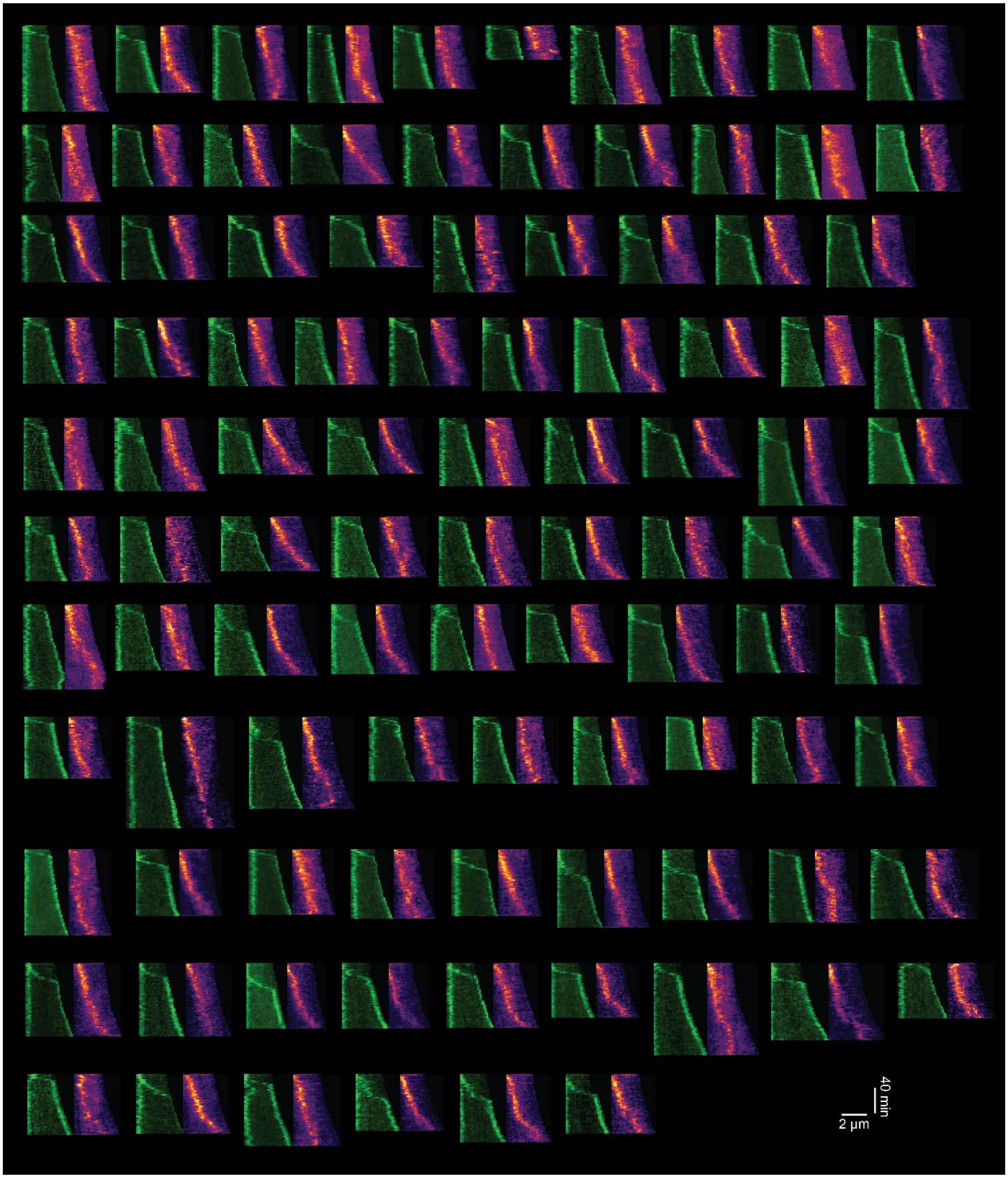


**Fig. S4:** Kymographs of ParB (left, green) and DnaN (right, pseudo-colored in MPI-inferno) fluorescence in WT (CB15N) background cells.

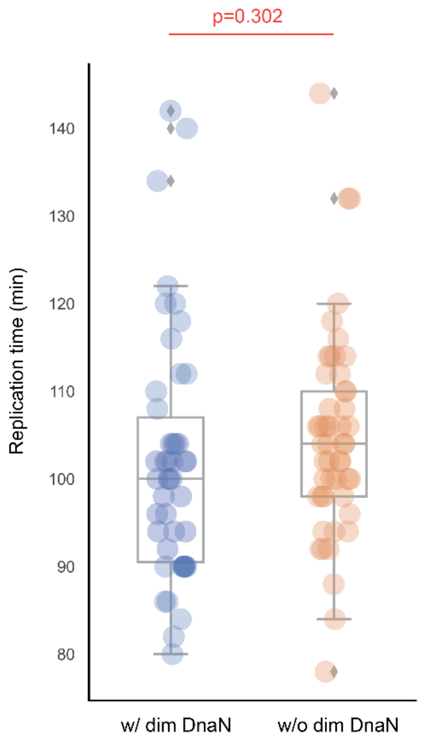


**Fig. S5:** Distribution of the replication time for cells with or without dim DnaN signals. The *p*-value is calculated by a two-tailed t-test.

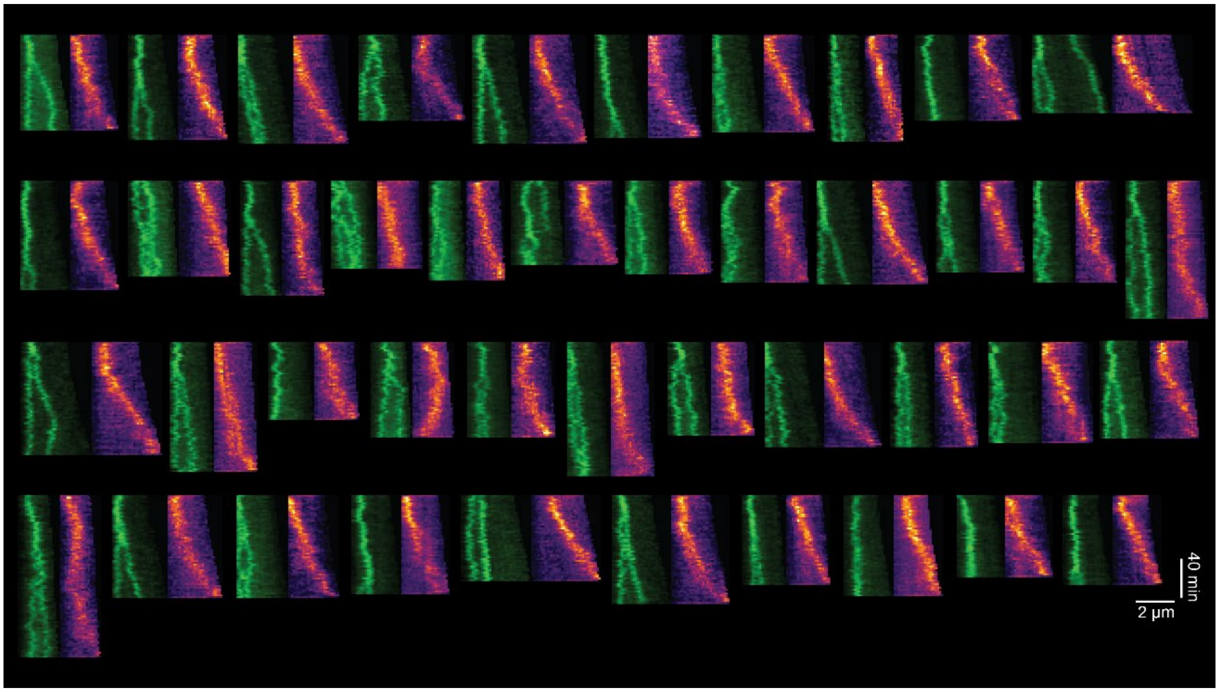


**Fig. S6:** Kymographs of ParB (left, green) and DnaN (right, pseudo-colored in MPI-inferno) fluorescence in *parAK20R* background cells.

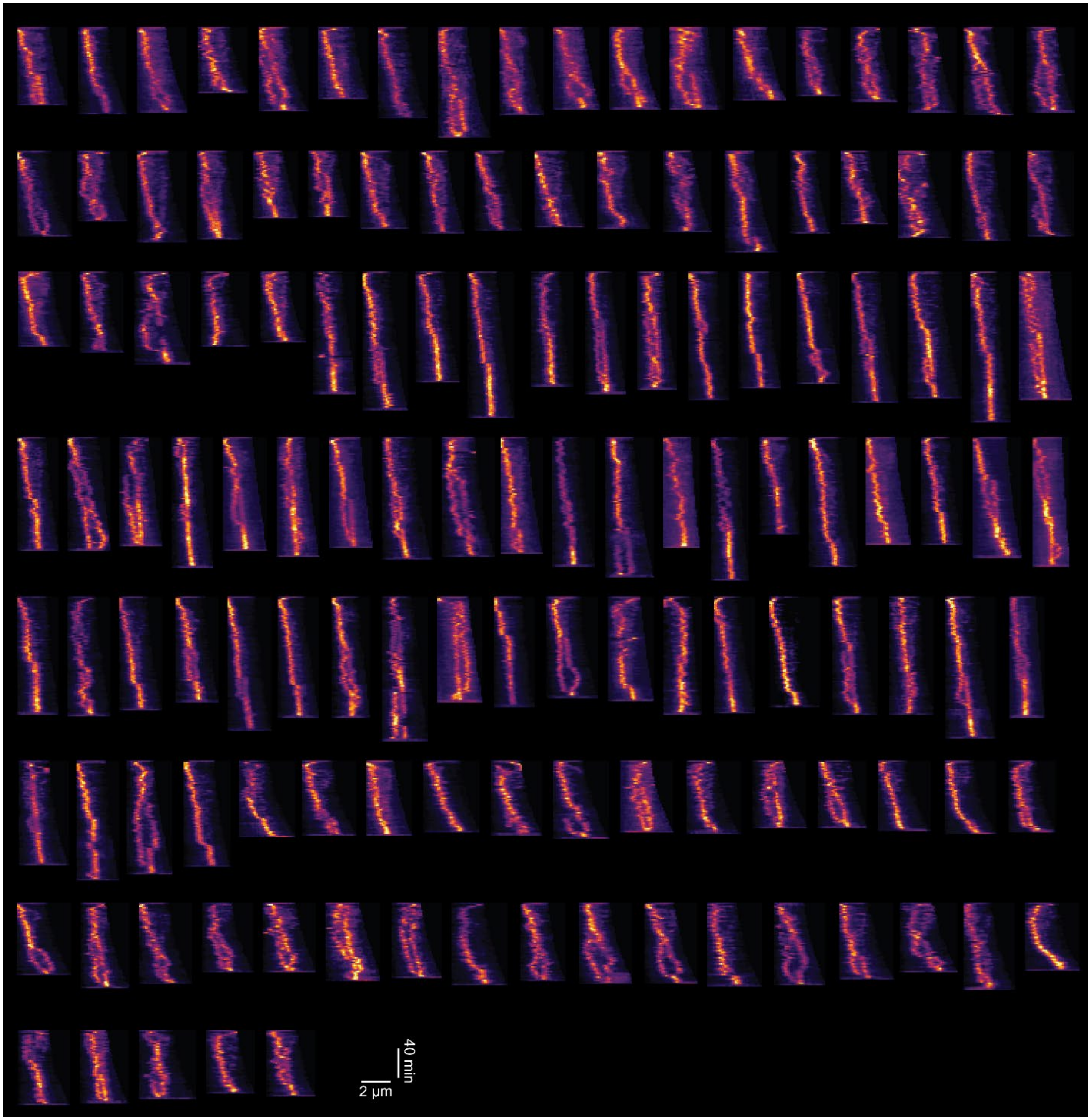


**Fig. S7:** Kymographs of time-lapse imaging of the CB15N Δ*smc::dnaN-sfGFP* cells.

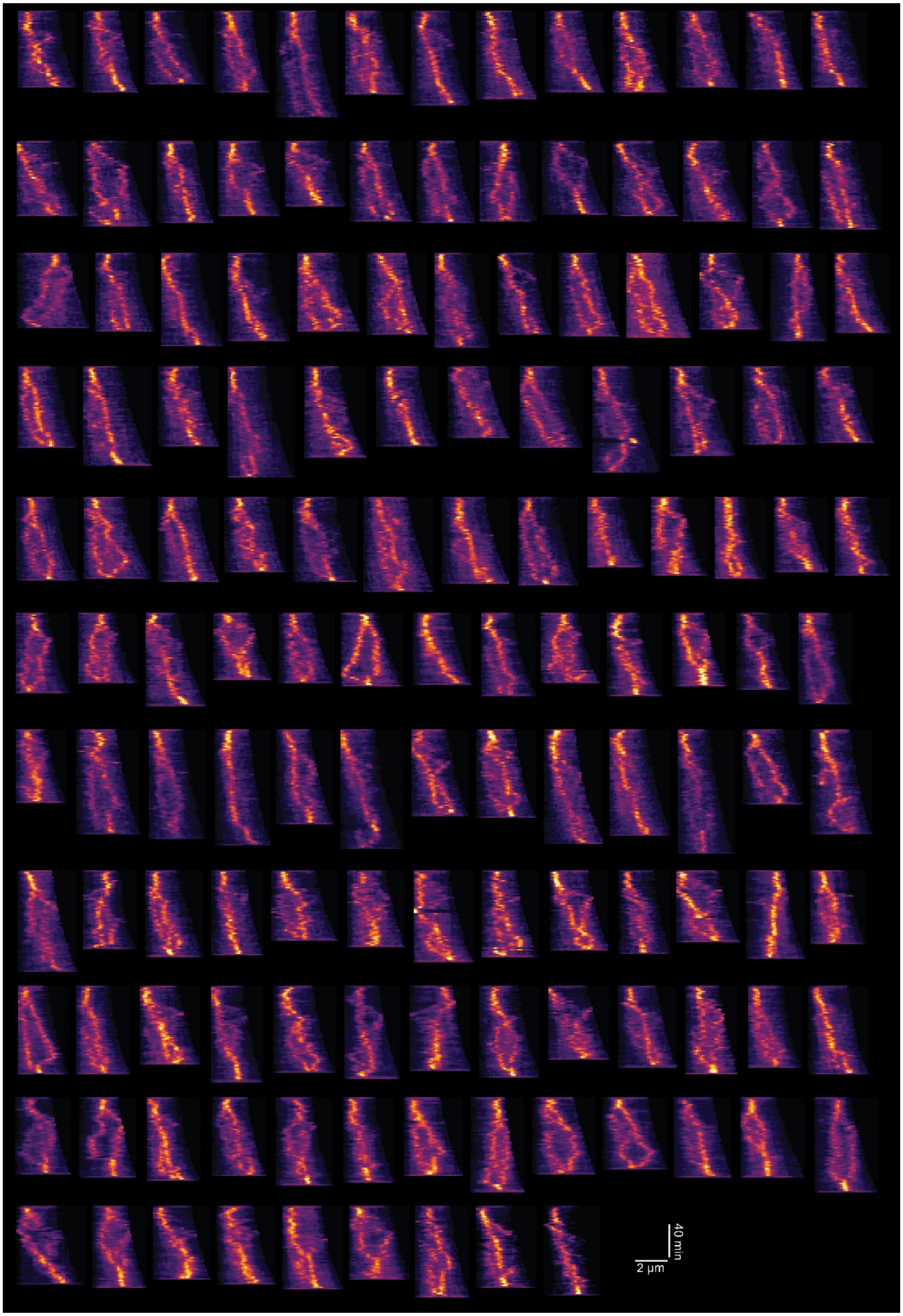


**Fig. S8:** Kymographs of time-lapse imaging of the CB15N *flip1-5::dnaN-sfGFP* cells.

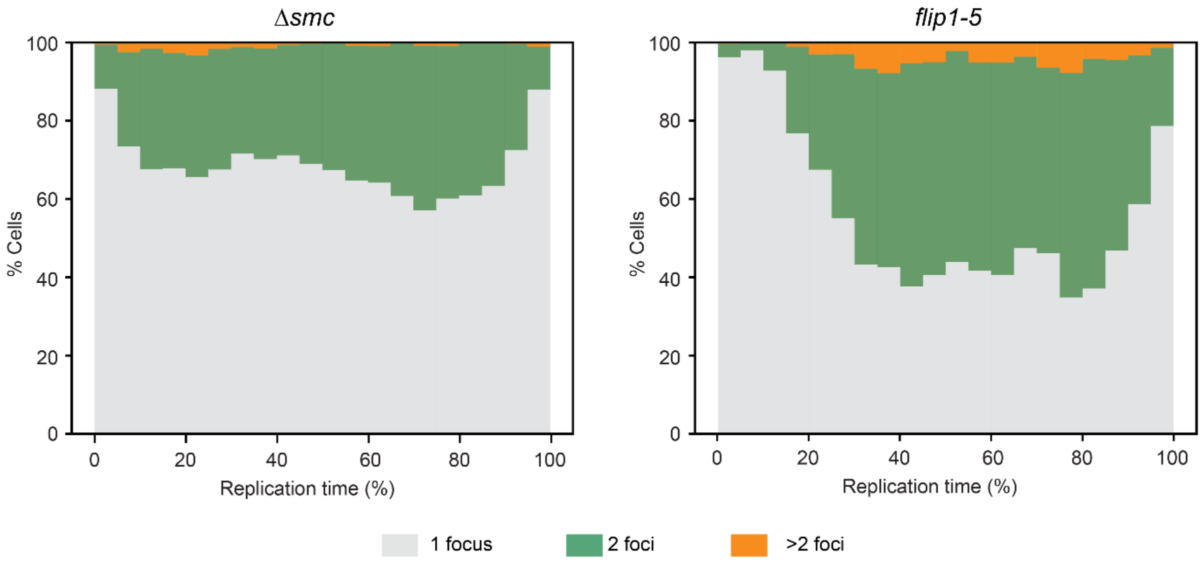


**Fig. S9:** Distribution of CB15N Δ*smc::dnaN-sfGFP* (left) and CB15N *flip1-5::dnaN-sfGFP* (right) cells that contain 1, 2, and >2 detected DnaN foci.

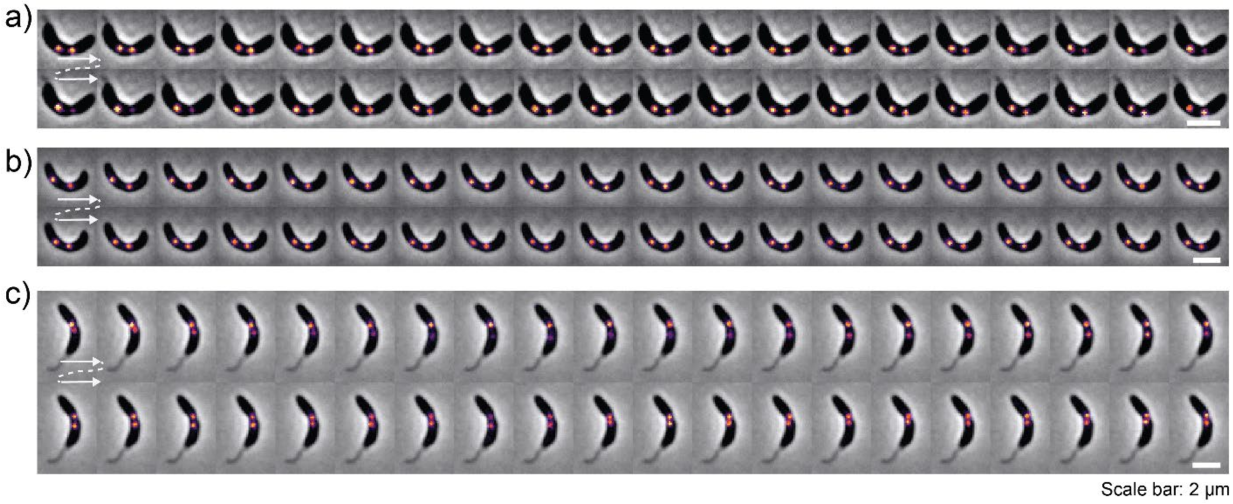


**Fig. S10:** Representative montages of DnaN fluorescence in pre-divisional CB15N*::dnaN-sfGFP* cells.

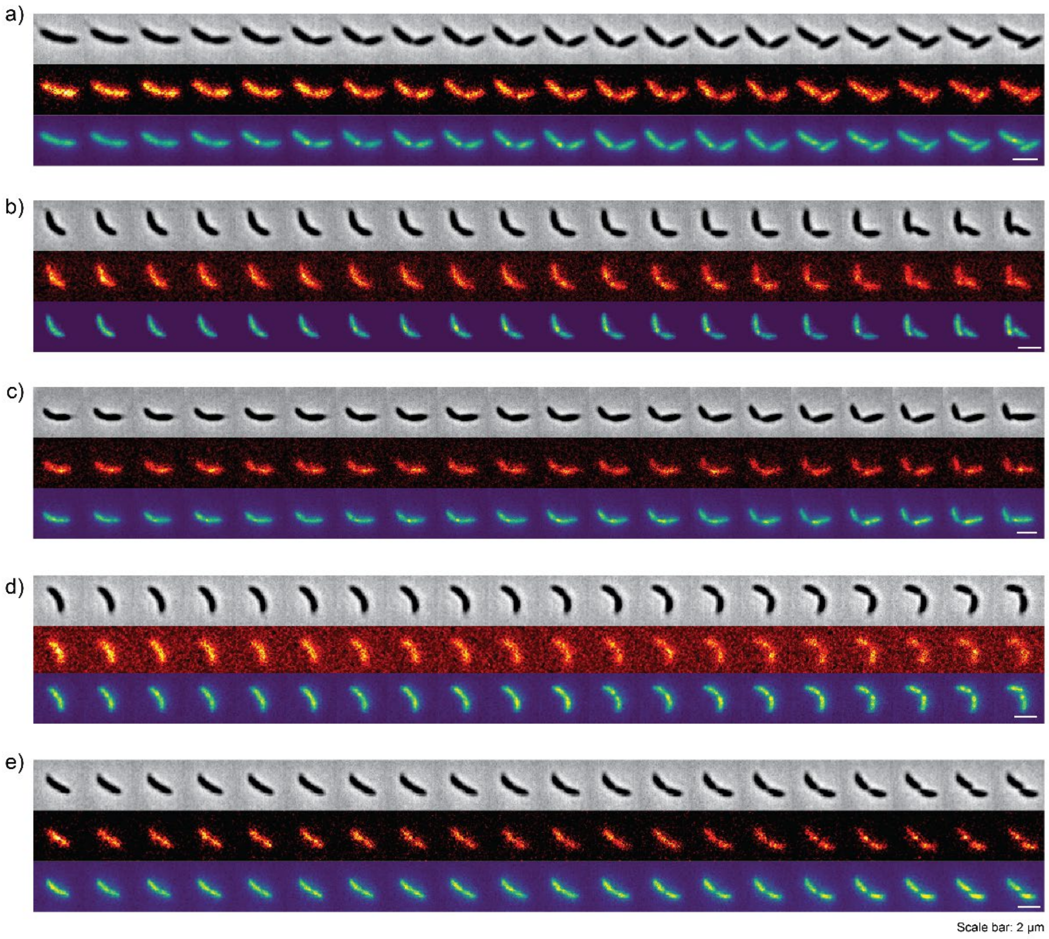


**Fig. S11:** Five example montages of phase contrast (top), and L5' (middle, pseudo-colored in Red-Hot), R5' (bottom, pseudo-colored in MPI-viridis) fluorescence of time-lapse imaging of CB15N*::L5'::R5'* cells with 3 min intervals.

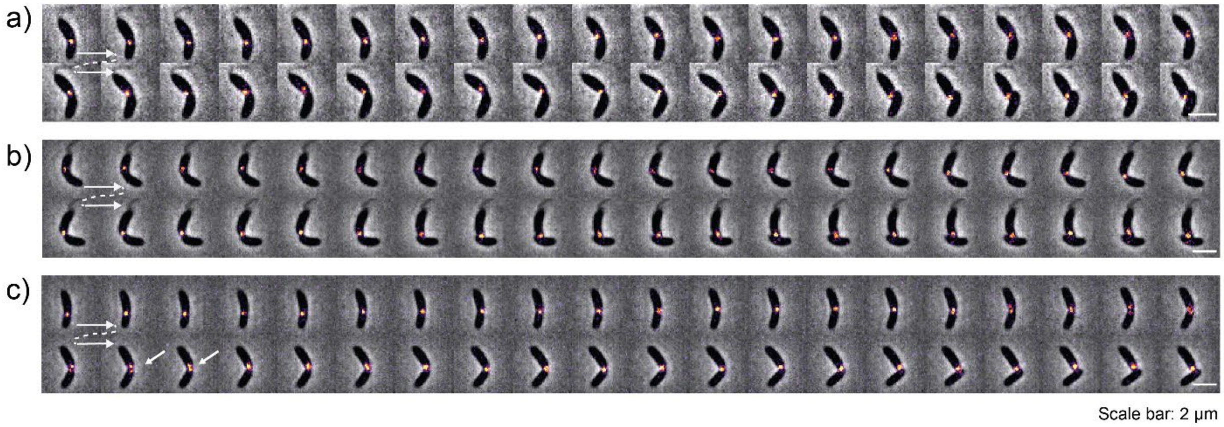


**Fig. S12:** Example montages showing the missing of loci splitting for CB15N*::R5* (a) and CB15N*::L5* (b), and the disappearance of split loci in CB15N*::R5* (c) with 2 min intervals.

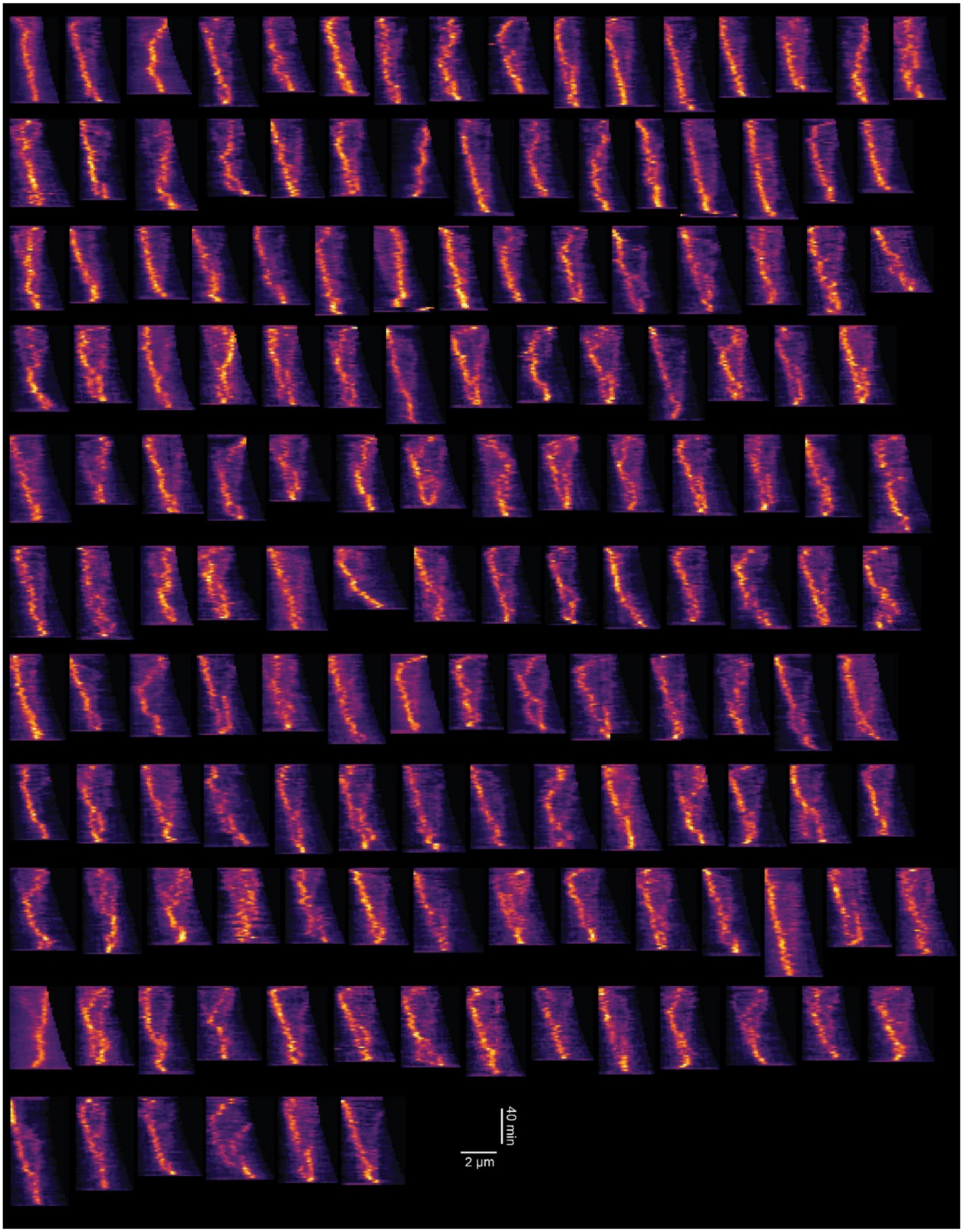


**Fig. S13:** Kymographs of time-lapse imaging of the CB15N Δ*rsaA::dnaN-sfGFP* cells.

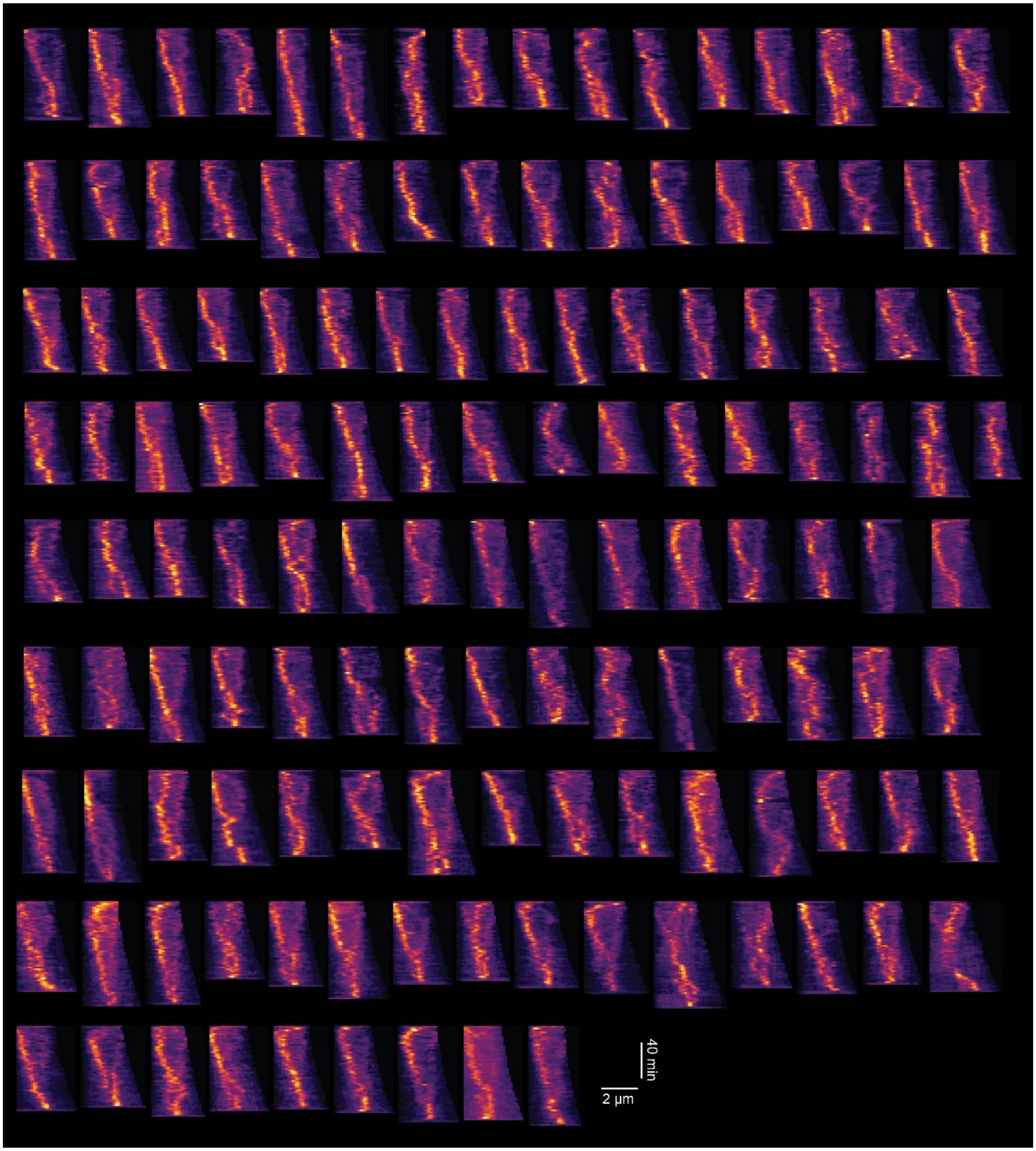


**Fig. S14:** Kymographs of time-lapse imaging of the CB15N Δ*rsaA::$P_{xyl}$-rasA::dnaN-sfGFP* cells.

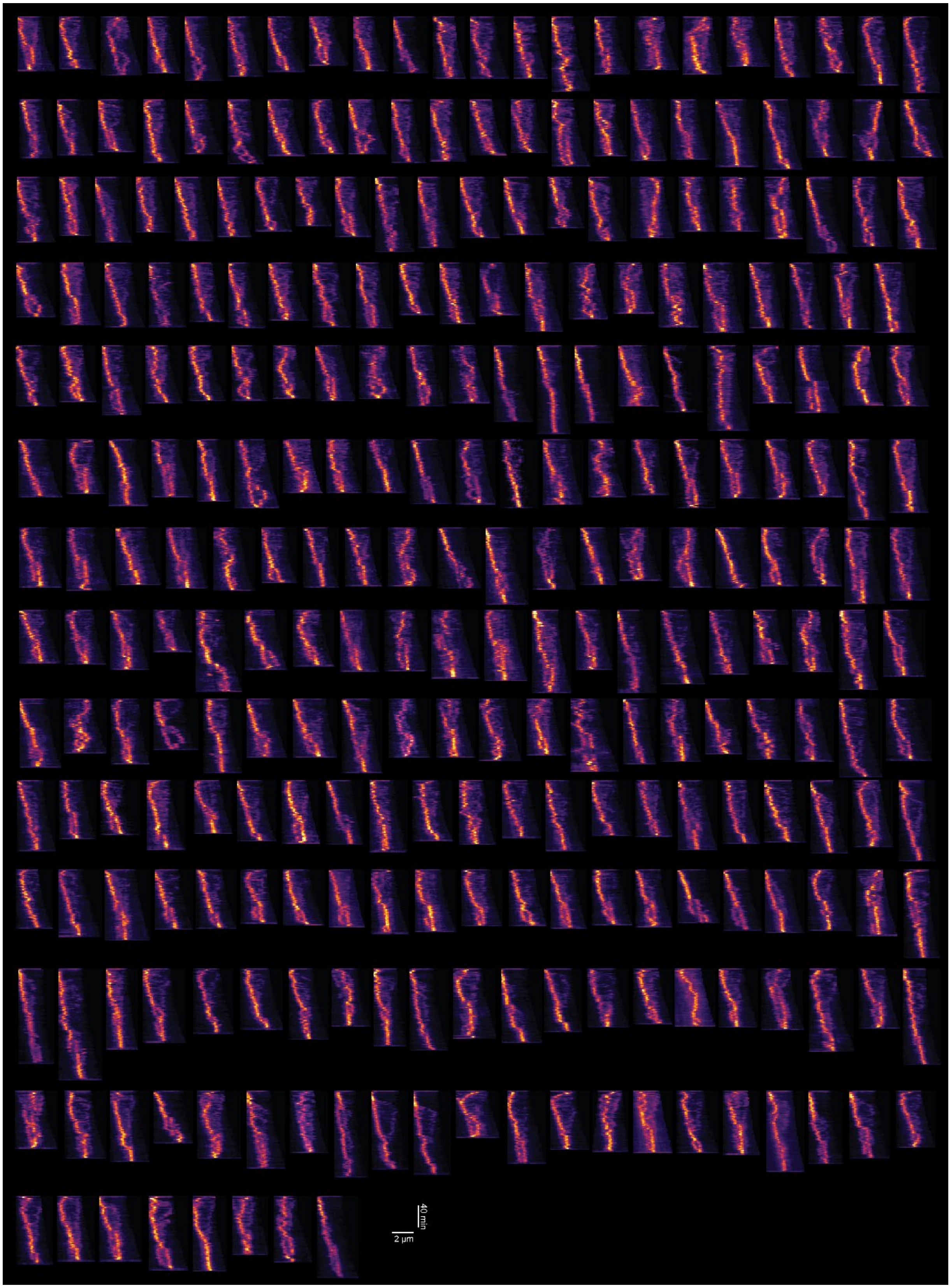


**Fig. S15:** Kymographs of time-lapse imaging of the CB15N*::P*$_{xyl}$*-rsaA::dnaN-sfGFP* (*rsaA*+) cells.

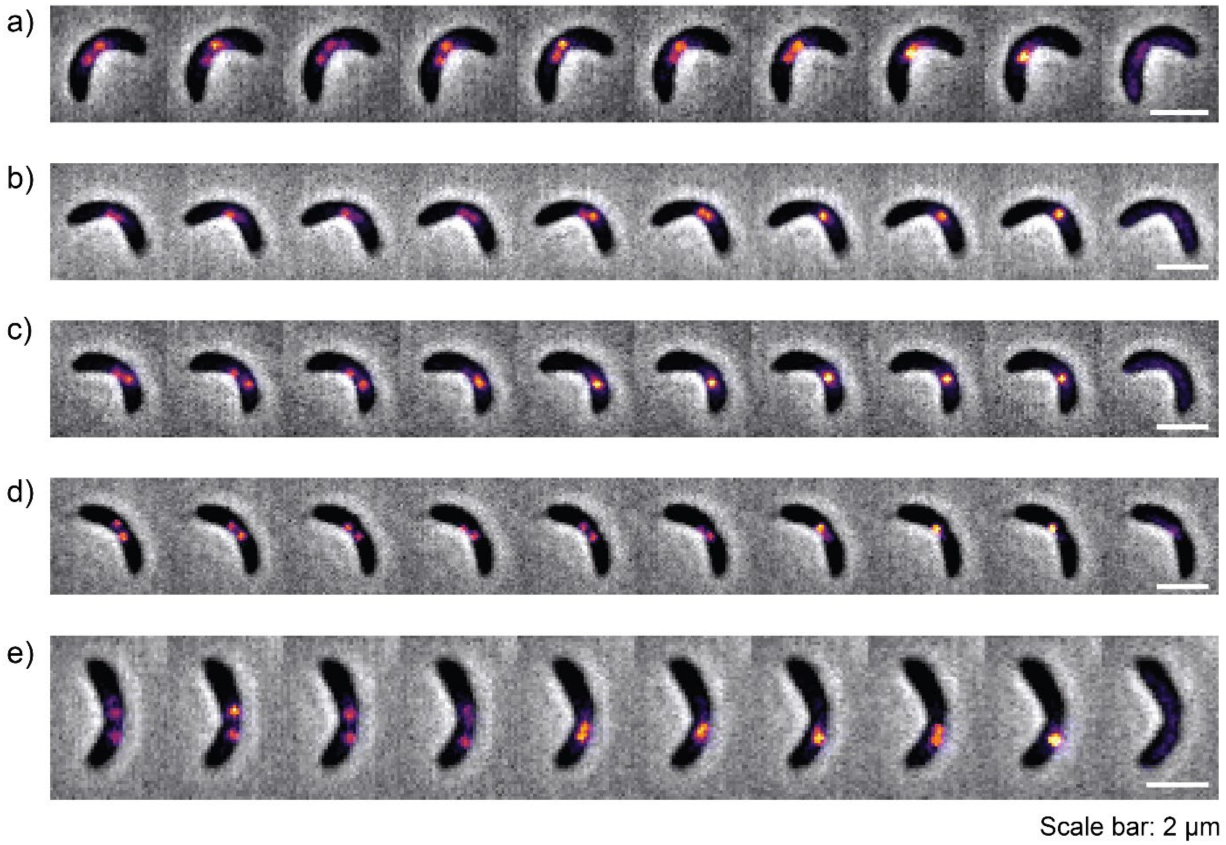


**Fig. S16:** Example montages of time-lapse contrast and DnaN fluorescence of *rsaA+* cells with a 2 min interval.

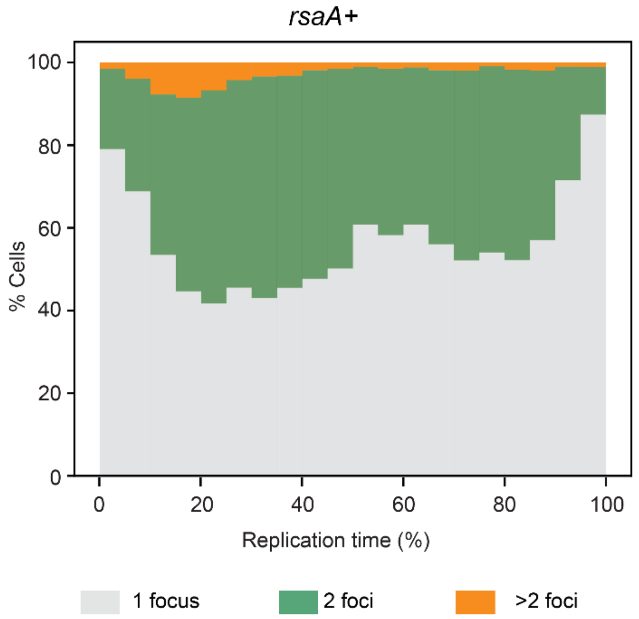


**Fig. S17:** Distribution of *rsaA+::dnaN-sfGFP* cells that contain 1, 2, and >2 detected DnaN foci.

**a**

Values used for estimating replication speed between genomic loci

| Loci | Loci position (kb) | Lengt at loci splitting (μm) |
|---|---|---|
| L2 | -371.0 | 2.51 |
| L3 | -987.3 | 2.68 |
| L4 | -1381.4 | 2.73 |
| L5 | -1521.7 | 2.88 |
| L6 | -1665.1 | 3.11 |
| R2 | 433.4 | 2.54 |
| R3 | 983.2 | 2.73 |
| R4 | 1375.8 | 3.00 |
| R5 | 1524.8 | 3.10 |
| R6 | 1687.3 | 3.39 |

An initial length L0 of 2.18 μm was assumed when replication starts.
To esitimat the replication speed on other loci, see example of R3 in **b**

**b**

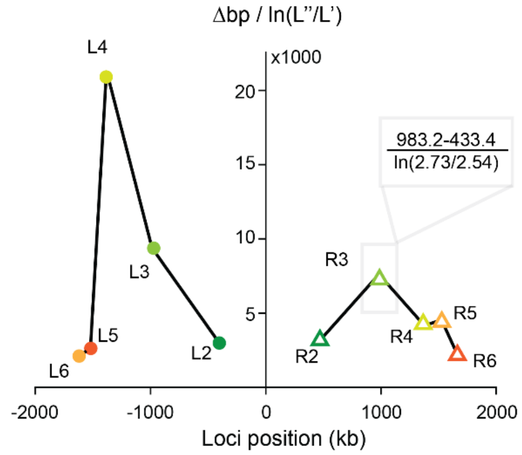


**Fig. S18**: (a) Values recorded for estimating the replication speed. (b) plot of the estimated speed at different genomic loci positions.

| | Doubling time (h) | | | | |
|---|---|---|---|---|---|
| | **WT** | **Δ*smc*** | ***flip1-5*** | **Δ*rsaA*** | ***rsaA+*** |
| M2G (28 °C) | 3.61 ± 0.14 | 3.49 ± 0.10 | 3.50 ± 0.06 | 3.61 ± 0.02 | 3.34 ± 0.06 |
| M2G (32 °C) | 2.96 ± 0.03 | 2.97 ± 0.05 | 2.93 ± 0.19 | 3.08 ± 0.10 | 2.79 ± 0.03 |
| PYE (32 °C) | 2.29 ± 0.11 | 2.26 ± 0.08 | 2.92 ± 0.05 | 2.33 ± 0.10 | 2.14 ± 0.09 |

**Fig. S19**: Doubling time of different background strains measured in nutrient and minimal medium by microplate reader. All strains harbor the same *dnaN-sfGFP* integration.
